# Effect of Wall Motion Sampling on CFD-Derived Left Atrial Flow Metrics

**DOI:** 10.1101/2025.10.24.684343

**Authors:** Yvonne Stöcker, Manuel Guerrero-Hurtado, Eduardo Durán, Alejandro Gonzalo, Zorica Ristić, Åshild Telle, Ahmad Kassar, Romanos Haykal, Nazem Akoum, Patrick M. Boyle, Oscar Flores, Christoph M. Augustin, Juan Carlos del Alamo, Manuel García-Villalba

## Abstract

The temporal resolution of medical imaging sequences used to drive patient-specific computational fluid dynamics (CFD) simulations remains limited, typically providing 10–20 frames per cardiac cycle. Therefore, temporal interpolation to reconstruct left atrial (LA) wall motion and boundary conditions is required, but its impact on hemodynamic predictions has not been systematically characterized. To investigate this, we constructed high-temporal-resolution reference wall-motion data using electromechanical (EM) simulations on five patient-specific atrial geometries with a history of atrial fibrillation. We then generated temporally downsampled datasets to emulate clinical frame rates (5, 10, 20, and 40 frames per cycle) and performed CFD simulations to isolate the effects of temporal undersampling on hemodynamic metrics. The focus was placed on kinetic energy, KE, and residence time, T*_R_*, particularly in the left atrial appendage (LAA), where thrombosis is most likely to occur. We employed an immersed boundary method to prescribe the wall motion and computed blood T*_R_* through a passive scalar transport equation.

Results indicate that while global LA hemodynamic indices were marginally affected by the frame rate (errors < 9%), LAA metrics were more sensitive with errors up to 31% compared to reference values. The results based on 20 and 40 frames per cycle yielded favorable agreement with reference results, while 5-and 10-frame reconstructions showed larger, though not systematically biased, deviations from the reference. Importantly, patient ranking by blood-stasis indices was largely preserved. The analysis suggests that patientspecific LA reconstructions derived from dynamic CT imaging provide a reliable basis for estimating LAA blood-stasis indices. Higher frame rates (*≥* 20 per cycle) offer improved quantitative accuracy, while lower temporal resolutions may remain informative for patient stratification purposes, where relative ranking is more relevant than absolute accuracy.

## 1 Introduction

Atrial fibrillation (AF) affects over 50 million people worldwide [1] and is characterized by weak, erratic contractions of the left atrium (LA). In AF, weakened LA motion promotes blood stasis, particularly in the left atrial appendage (LAA)—a small, narrow-necked chamber whose emptying is especially compromised under reduced contractility. This predisposes to thrombus formation in the LAA, which can embolize and cause ischemic stroke [2]. Anticoagulation therapy reduces stroke risk but increases bleeding risk, making risk stratification essential to ensure a net clinical benefit [3, 4]. Commonly, stroke risk stratification relies on scores such as the CHADS_2_ score and its variations, which are purely based on patient demographics and medical history factors derived from large population-based studies [5]. Notably, they lack information about patient-specific hemodynamics [6, 7], and their predictive accuracy is moderate [8].

Because blood stasis is a major thrombogenic factor [9], patient-specific computational fluid dynamics (CFD) analyses hold promise for improving individualized stroke risk estimation [10–13]. Accurate modeling of LA hemodynamics is essential to predict thrombus formation, and recent years have seen substantial progress in this area [14]. However, CFD results can be sensitive to modeling assumptions and numerical methods. Some studies have analyzed the effects of spatiotemporal resolution and numerical order of discretization [15], or the potential use of turbulence models in LA simulations [16], where flow remains predominantly laminar during most of the cardiac cycle. Others have assessed the effects of non-Newtonian rheology of slow-flowing blood [12, 17], inflow boundary condition choices [18], including the pulmonary vein (PV) flow split [16, 19–21], and presence of the mitral valve (MV) [22].

Constructing patient-specific CFD models requires segmenting the interface between atrial tissue and blood—hereafter, the LA *wall* —from 3D medical images and generating computational meshes. Capturing wall motion further requires time-resolved imaging (i.e., 4D), frame-by-frame segmentation, and registration across frames over the cardiac cycle. However, 4D cardiac imaging is not routinely performed at many centers. Moreover, in patients with paroxysmal AF, the rhythm present at the scan cannot be controlled; patients may be imaged in AF or in sinus rhythm, which further complicates model construction. These constraints have led many investigators to use rigid-wall simulations to approximate impaired wall motion during AF [10, 12, 23–29], or even sinus rhythm [10, 15, 16, 25, 30]. Several studies have evaluated this approximation [16, 18, 31, 32], concluding that it yields unphysiological flow in sinus rhythm whereas it tends to overestimate thrombosis predictors in AF.

Motivated by these limitations, researchers have utilized simplified motion laws [6, 33–38], fluid–structure-interaction approaches [39–45], prescribed wall motion obtained from prior electromechanics (EM) simulations [46, 47], or applied deep learning approaches to recreate wall motion from still images [48]. Although more sophisticated than assuming rigid LA walls, these strategies still depend on modeling choices and require careful parameterization or training. Consequently, when available, CFD driven by 4D medical images (typically magnetic resonance imaging (MRI) or computed tomography (CT)) is commonly adopted [17, 20, 22, 30, 31, 49–59]. However, these modalities provide limited temporal resolution (*∼* 10 frames per cycle), necessitating temporal interpolation, and the impact of this undersampling on simulation-derived metrics has not been systematically characterized.

In this work, we quantify how time resolution of wall motion affects atrial hemodynamics in CFD simulations. To address the lack of high-frame-rate image sequences to build ground-truth LA models, we construct a high-temporal-resolution reference of wall motion utilizing EM simulations on five patient-specific atrial geometries. Subsequently, we generate temporally downsampled datasets that emulate clinical frame rates and perform new CFD simulations. In this way, we isolate the effect of temporal undersampling on hemodynamic metrics.

In the following section, we present the methodology of the EM and CFD simulations, including the wall-motion interpolation strategy. In section 3, we first give an overview of the EM results, followed by detailed results of the CFD simulations for ground-truth and undersampled wall-motion cases, with particular emphasis on the LAA, the most common site of atrial thrombosis in AF. In the final part of section 3, we investigate the effects of downsampling influencing not only wall motion but also the prescribed transmitral outflow rate. Section 4 discusses these results and addresses the study limitations, while section 5 summarizes the main results of this study.

## 2 Methodology

### 2.1 Ethics statement

The study (STUDY00015081) was approved by the Institutional Review Board of the University of Washington. All patients provided their written consent.

### 2.2 Patients & Imaging

The five patients considered here were recruited from the University of Washington Medical Center, three of whom (B, D, and E) are the same as in Telle *et al.* [60]. They all had a history of AF and were scheduled for ablation. MRI scans were taken prior to ablation at the end of atrial diastole and then segmented, processed, and analyzed by Merisight (Marrek Inc., Salt Lake City, UT) as described in Marrouche *et al.* [61].

### 2.3 Mesh generation

To perform EM simulations, the LA myocardium was meshed using *meshtool* [62], an open-source software for mesh generation and manipulation tailored to cardiac modeling applications. First, the interface between blood pool and tissue was extracted from the image segmentations and discretized with triangular elements, yielding the surface meshes displayed in Figure 1.

**Figure 1:**
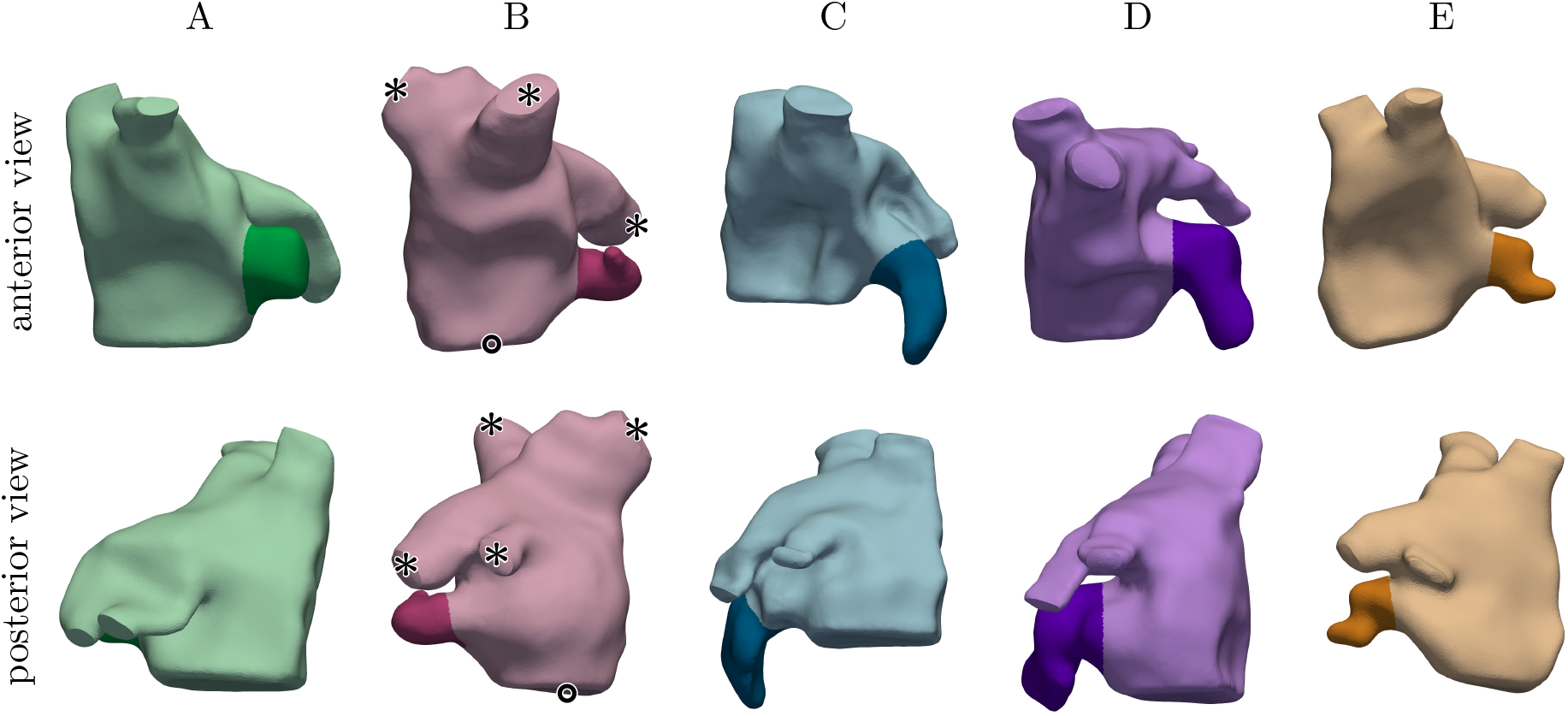
Left atrial anatomies of the five study subjects, showing individual color assignments and delineation of the LAA in each case. For subject B, the locations of the PVs are indicated by an asterisk, and the MV by a circle.

Subsequently, a volumetric, tetrahedral mesh of the myocardial tissue was generated by extruding the blood pool interface by 0.8 mm to generate the endocardium and by a further 1.2 mm to create the epicardial layer, resulting in an overall atrial wall thickness of 2 mm [63]. Atrial myocyte orientations were assigned based on automatically identified anatomical landmark regions, following previous works [64, 65]. The average tetrahedral edge length was approximately 500 µm.

Additionally, auxiliary volumetric patches were introduced to close the MV and PV orifices. These caps facilitate the definition of boundary conditions in both the EM and CFD simulations, and help to preserve the anatomical shape of the orifices [60].

### 2.4 EM simulations

Based on the patient-specific anatomies, myocardial EM simulations were performed using the in-house code *CARPentry* [66, 67]. The human atrial action potential was represented by the Courtemanche *et al.* [68] model of cellular electrophysiology, with modifications as described in Bayer *et al.* [69]. Excitation of the LA was initiated through transmembrane current stimulation in three selected points around the right superior PV, anatomically representing approximate activation sites as observed for patients B, D, and E in a previous study [60]. The propagation of electrical activation in the myocardium was simulated using the reaction– eikonal equation [70], which enables consistent and physiological activation patterns regardless of mesh discretization. Longitudinal and transverse conduction velocity was set to 1.308 m/s and 0.585 m/s, respectively, based on average values reported for patients B, D, and E [60]. Active stress generation was represented by the model of human atrial contraction dynamics described in Land and Niederer [71].

Passive tension in the LA was described using a reduced Holzapfel–Ogden formulation with fiber dispersion [72], while the auxiliary PV and MV caps were represented by a stiff Demiray material [60]. Both materials were treated as nearly incompressible. Robin-type boundary conditions were used to constrain atrial motion: normal springs on the epicardium and omnidirectional springs at the PV inlets. For more information on the EM model, including model parameters, see [46, 60, 72].

Since the exact pressure at the time of image acquisition was unknown, an unloaded reference configuration was estimated using a backward displacement algorithm [73] with a predefined pressure of 10 mmHg. This procedure ensures that the pressurized anatomical model matches the geometry obtained from imaging data.

To account for the hemodynamic interactions with the remaining parts of the circulatory system, the 3D EM model of the LA was strongly coupled to *CircAdapt* [74, 75], a non-linear 0D lumped parameter model that efficiently simulates the dynamics of the other heart chambers as well as the systemic and pulmonary circulation. *CircAdapt* simulated hydrostatic pressures and blood flow across the PV inlets and MV outlet, which were used to impose physiological pressure boundary conditions and volume constraints on the solid LA structure [67]. Additionally, the ventricular contraction in the 0D model was used to approximate atrioventricular plane displacement, which was applied as a traction boundary condition on the MV annulus following Telle *et al.* [60].

Electrical activation of the LA was initialized at 0 ms of each heartbeat at a heart rate of 60 bpm. To ensure periodicity, 28 beats were first simulated using a coarser mechanical time step of Δt_mech_ = 1 ms, followed by two additional beats with a refined time step of Δt_mech_ = 50 µs, consistent with the requirements of the CFD simulations described below. For the electrophysiology components, a uniform time step size of Δt_EP_ = 25 µs was used throughout all beats. The final state from the 30^th^ beat was then provided to the CFD pipeline.

### 2.5 CFD simulations

Assuming incompressibility and Newtonian blood rheology, the Navier–Stokes equations that govern the dynamics of the blood flow were solved with a fractional step method using the inhouse code *TUCANGPU* [76], which is the GPU-accelerated successor of the CPU-based flow solver *TUCAN* [77]. The same heart rate (60 bpm) and blood viscosity (0.04 cm^2^/s) were applied across all subjects to standardize comparisons. Discretization in space was performed with second-order central finite differences in a staggered, uniform Cartesian grid. For integration in time, a low-storage, semi-implicit Runge–Kutta scheme with three stages was employed.

Simultaneously, the blood residence time T*_R_* was computed from the forced transport equation for a passive scalar

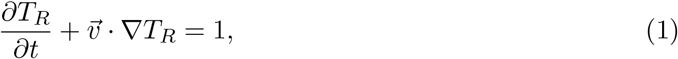

where v⃗ is the fluid velocity vector. This equation was spatially discretized with a third-order WENO scheme [78] to ensure accuracy and avoid spurious oscillations near sharp gradients.

Each simulation was performed in a triply-periodic cubic computational domain with an edge length of 13 cm and 256 uniformly distributed points per direction, resulting in a grid step size of approximately Δx = 0.51 mm (16.8 million grid cells). The time step size Δt_CFD_ = 50 µs was kept constant and chosen to achieve a Courant number CFL < 0.3 throughout the simulations.

To couple the flow to the motion of the LA wall, an immersed boundary method (IBM) [79] was used. In this approach, the moving LA wall is placed in the fixed computational domain and the no-slip condition at the LA surface is imposed by adding appropriate localized volumetric forces as a source term in the momentum equations of the fluid. These IBM forces are evaluated at the location of specific marker points, which are distributed over the LA surface. They correspond to the center points of the triangular surface elements of the inner myocardial wall (also including the PV and MV caps), whose motion is imposed as presented in the next section. To reduce leakage across the immersed boundary, which has also been observed by other groups [80, 81], a second layer of marker points was added by shifting each original marker by one grid step size Δx along the outward normal direction of its corresponding triangular element.

The flow rate through the PVs was enforced via a prescribed surface-normal velocity in a non-circular buffer region upstream of each PV orifice cap. It is computed based on mass conservation

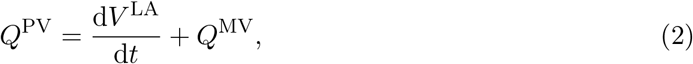

where V ^LA^ is the LA volume and Q^MV^ the known transmitral outflow rate obtained from the EM simulations. This Q^PV^ is then split between all PVs such that equal velocity is obtained in each vein. The buffer region at the PV inlets is also used to set the residence time T*_R_* to zero.

When outflow is desired, i.e. Q^MV^ > 0, no IBM force is imposed at the marker points that belong to the MV orifice cap. Otherwise, a no-slip condition is imposed. No specific model is applied for the MV leaflets, since preliminary studies showed no substantial differences in terms of the results presented in section 3.

All simulations were initialized with zero velocity and residence time, and run for 20 cardiac cycles to ensure convergence to a quasi-periodic state. This long convergence time corresponds to the washout time of the fluid particle with the highest residence time, which, especially in the LAA, can be very high. Results were collected from the subsequent eight cycles.

### 2.6 Wall motion

In previous atrial flow studies of our group, the LA geometry and its motion were obtained from patient-specific time-resolved computed tomography (4D–CT) images [17, 20, 31], which had a temporal resolution in the order of 10 to 20 images per cardiac cycle. However, with the duration of a cardiac cycle set to T = 1 s in the CFD simulations, and as they are carried out with a time step of 50 µs, motion information at 20 000 time instants per cycle is required. To achieve this, Fourier interpolation in time was applied to the marker point coordinates, with the number of Fourier modes depending on the number of available CT frames.

In contrast to the 4D–CT scans, EM simulations can supply such high temporal resolution that no interpolation would be needed. This provided the novel opportunity to assess how sensitive the CFD results are to the motion interpolation by comparing CFD results based on low-resolution, interpolated motion with results from fully resolved motion.

First, a reference simulation was performed for each subject by prescribing the LA wall motion from the full-resolution motion data from the EM simulations, i.e. without applying any interpolation. Then, the temporal resolution of the motion data was artificially decreased by retaining only a predefined number n*_f_* of equidistant time frames of one cycle, thereby mimicking the data available from 4D–CT scans. Subsequently, the motion was reconstructed using Fourier interpolation. The number of retained frames was chosen to be lower (n*_f_* = 5), equal (n*_f_* = 10 and 20), and higher (n*_f_* = 40) than in previous studies [17, 20, 31, 49, 50, 52, 58]. In the following, these simulated cases will be referred to by a letter identifying the subject, followed by a case identifier that denotes the number of retained frames n*_f_*. E.g., the cases of subject A are named Aref, A5, A10, A20, and A40.

## 3 Results

This study utilized patient-specific EM and CFD LA models from N = 5 subjects obtained from MRI. Table 1 summarizes relevant model-predicted anatomical and global functional chamber indices for each subject, as well as time-averaged LA residence time *T*_*R*_^LA^. The table includes a queue-model estimate for *T*_*R*_^LA^ based on global indices previously shown to approximate reference values from patient-specific CFD simulations based on 4D–CT segmentations [17, 20] (see Martínez-Legazpi *et al.* [82] for further details about cardiac chamber queue models).

**Table 1:** Model-predicted anatomical and functional parameters of the LA and LAA. Mean values represent time-averaged quantities. EF: emptying fraction, LVSV: left ventricular stroke volume. *T*_*R*_^LA^ represents residence time in the LA. A conduit-dominated queue model estimate (top) is 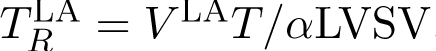 = V ^LA^T/αLVSV, where T is the cardiac cycle and the parameter α = 1.4 is fixed fitting previous data [20]. The colors are used to identify the subjects throughout the manuscript figures.

| subject |  | A | B | C | D | E |
| --- | --- | --- | --- | --- | --- | --- |
| mean $V$ [mL] | LA | 101 | 122 | 134 | 130 | 66 |
|  | LAA | 4.8 | 4.0 | 12.0 | 10.2 | 2.0 |
| EF $\left[ \frac{V_{\max} - V_{\min}}{V_{\max}} \right]$ | LA | 0.43 | 0.36 | 0.35 | 0.36 | 0.56 |
|  | LAA | 0.44 | 0.45 | 0.40 | 0.41 | 0.49 |
| LVSV [mL] |  | 85.6 | 83.2 | 83.7 | 84.5 | 89.3 |
| mean $T_R^{\text{LA}}$ [s] | estimated | 0.84 | 1.05 | 1.14 | 1.10 | 0.53 |
|  | CFD, ref | 1.07 | 1.05 | 1.46 | 1.50 | 0.62 |

While the functional parameters are similar across the subjects, the mean volumes vary significantly. In particular, patient E has markedly smaller LA and LAA volumes than the other subjects, resulting in a considerably lower residence time, as will be shown below. The LA and LAA emptying fractions are on the low end of normality in healthy adults [83], helping to set a more stringent benchmark for wall-motion models than previous studies that considered weakly beating atria [32].

The remainder of this section reports detailed results for subjects A and D, followed by statistical quantities for all five subjects. Particular attention is given to flow specific kinetic energy KE = *|*v⃗*|*^2^/2 and residence time (see Equation 1) as hemodynamic metrics, and on the LAA, the most likely thrombosis region. Results are collected at 40 equidistant time instants per cardiac cycle in the 21^st^ to 28^th^ cycle, and are obtained by considering only the Eulerian voxels that are inside the respective regions of interest.

### 3.1 Electromechanical simulations results

Representative results of the EM simulations for subjects A and D are shown in Figure 2, taken from the final, 30^th^, simulated cycle, after the solution had converged. The earliest activation sites, represented in dark blue, correspond to those identified in the electro–anatomical maps. Electrical activation then propagates throughout the remaining tissue, triggering active tension generation (Fig. 2b). This leads to active contraction in the cells and thus to deformation of the tissue, which in turn produces both passive and active mechanical stress (Fig. 2c). Finally, the interaction with the 0D model of the circulatory system yields the physiological pressure–volume relationship depicted in Figure 2d.

**Figure 2:**
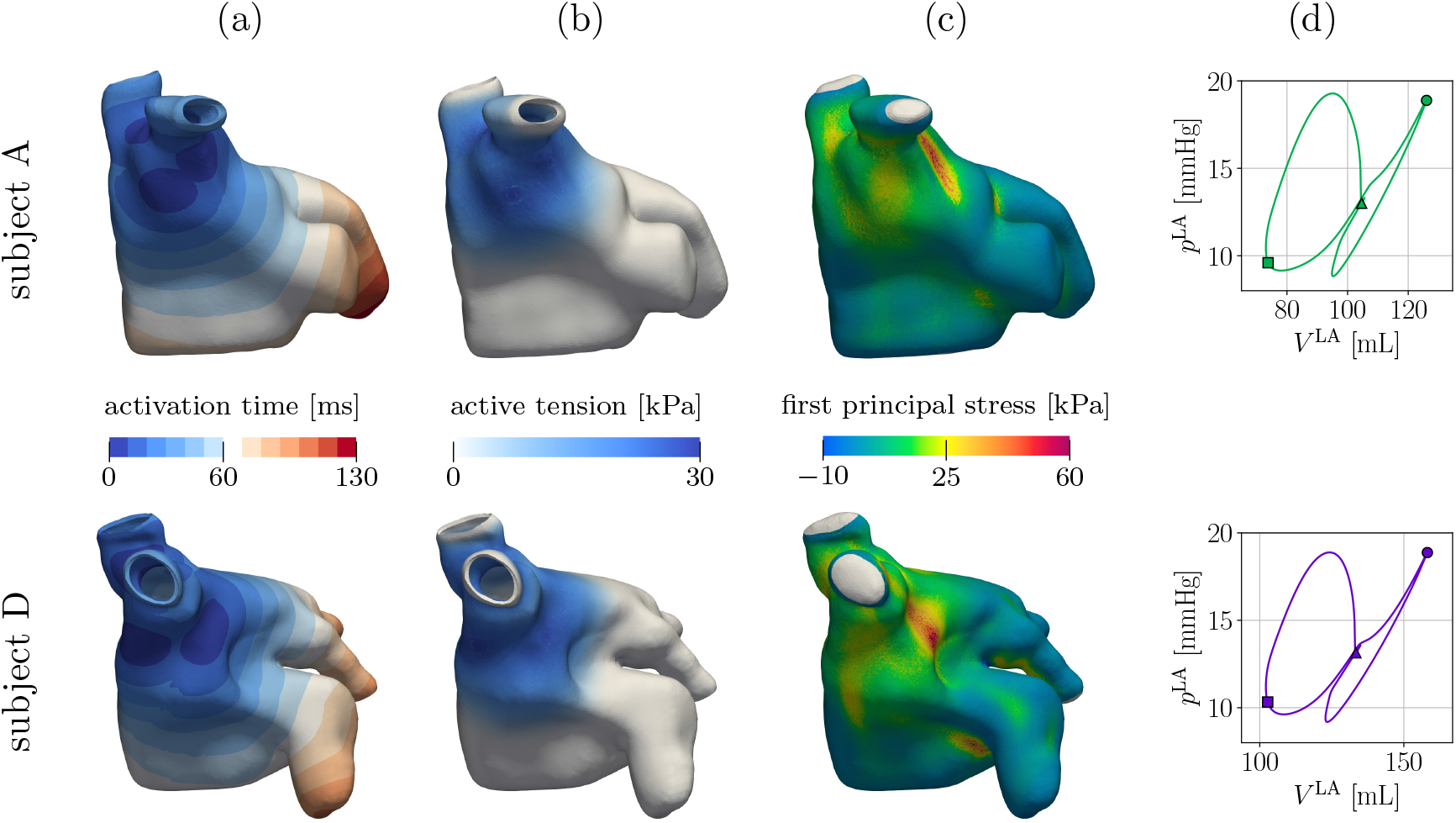
(a) Electrical activation sequence with earliest activation sites shown in dark blue. (b, c) Instantaneous spatial distribution of model outputs at 60 ms after atrial excitation: (b) active cellular tension and (c) first principal component of the total mechanical stress (sum of active and passive stress). (d) Left atrial pressure–volume curve with markers indicating the • MV opening, ■ MV closure, and ▲ point corresponding to panels (b, c). In (c), the PV caps are shown in gray to indicate that stress distribution there is not relevant for further analysis. Since they are modeled as non-conductive passive tissue, they are omitted in (a, b).

### 3.2 Impact of LA Wall-Motion Temporal Resolution on Global Blood Mass Conservation

We first assess how the temporal resolution of the LA wall reconstruction and its interpolation affect global mass conservation for the fluid inside the chamber, since mass balance (Eq. 2) is used to prescribe PV inflow boundary conditions. Figure 3a,b displays the temporal derivative of LA volume dV ^LA^/dt, the MV outflow rate Q^MV^, and the PV flow rate Q^PV^ over time for one cycle of the reference cases Aref and Dref. Atrial contraction (A-wave) occurs at t/T *≈* 0.1, followed by atrial expansion and the left ventricular filling (E-wave), which starts at approximately t/T *≈* 0.55.

**Figure 3:**
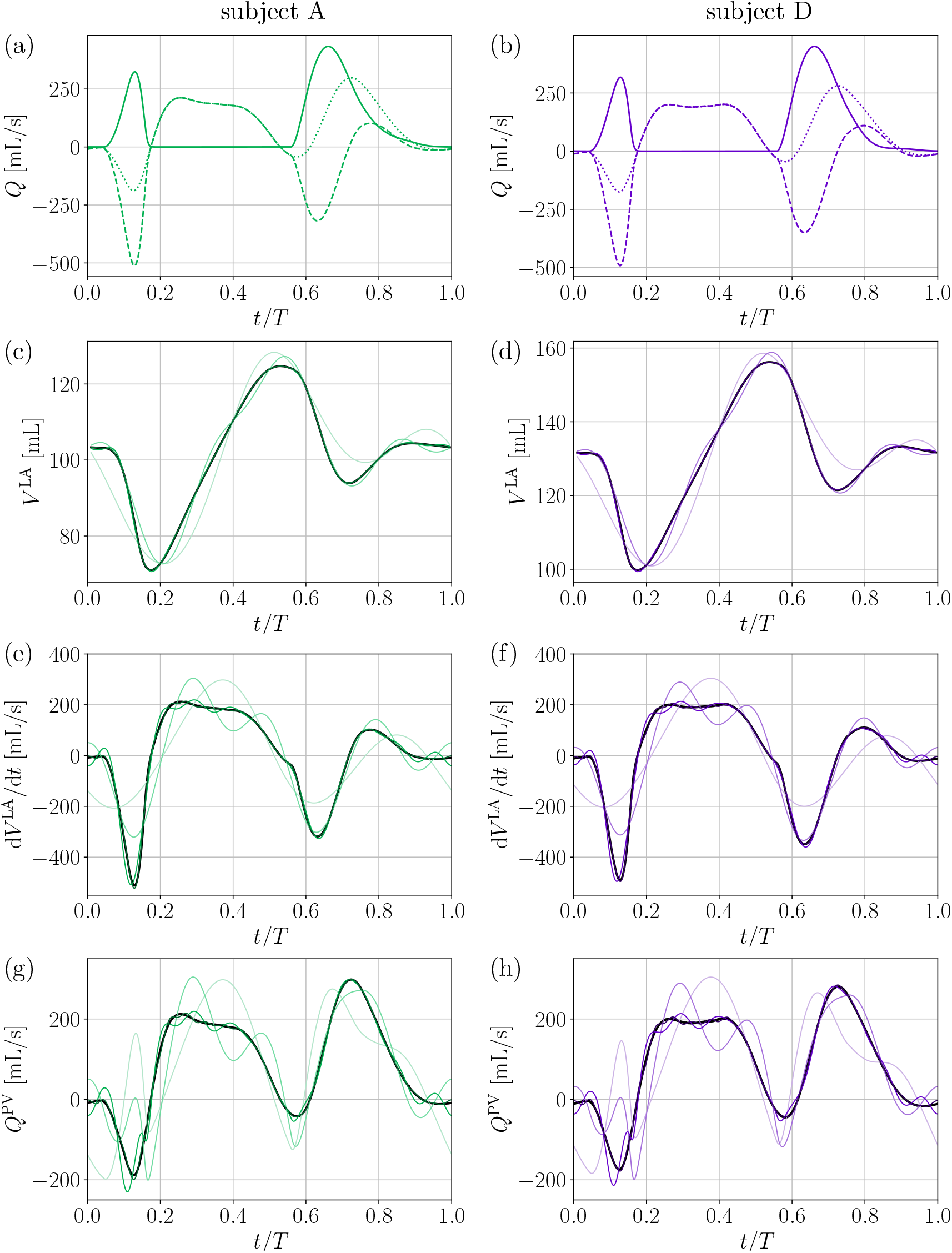
Temporal evolution of flow quantities during one cycle for subjects (left) A and (right) D. Q^MV^, Q^PV^, and dV ^LA^/dt in (a) Aref and (b) Dref; (c, d) Temporal evolution of LA volume, (e, f) time derivative of LA volume, and (g, h) PV flow rate. In (c–h), the 5-, 10-, 20-, and 40-frame reconstructions are plotted in color scheme with an increasing number of frames from lighter to darker color, while the reference 4D LA reconstruction is plotted in black.

Resulting from the interpolation, the evolution of LA volume in the reconstructed cases differs from their respective reference cases, which are depicted in Figure 3c,d. The reconstructed cases coincide with the reference ones only at the time instants of sampling. Consequently, as n*_f_* increases, the curves approach their reference graphs, with the 40-frame cases producing almost a perfect reconstruction.

In Figure 3e,f, the temporal derivative of LA volume is shown for all five cases of subjects A and D. In the interpolated reconstructions, the curves exhibit Fourier-induced spurious oscillations, whose amplitude grows and frequency falls as the number of retained 4D frames decreases. Notably, key physiological landmarks, e.g., the E-and A-wave peaks, are smeared and attenuated in the interpolated reconstructions with n*_f_ ≤* 10 retained frames.

As anticipated, these differences in dV ^LA^/dt could impact the CFD simulations because the outflow rate through the MV is prescribed identically for all cases of the same subject, regardless of the temporal resolution of its 4D LA wall-motion model. Thus, since the flow through the

PVs is prescribed to preserve mass while enforcing the target outflow (Eq. 2), reconstruction-dependent differences in dV ^LA^/dt can produce significantly different inflow boundary conditions Q^PV^ within the same subject, as shown in Figure 3g,h. The mean absolute errors made in Q^PV^ of the reconstructions with respect to the reference flow rate are summarized in Table 2, showing that this error systematically increased as n*_f_* was lowered. The next section evaluates how these differences affect main hemodynamic metrics such as KE and T*_R_*.

**Table 2:** Mean absolute error [mL/s] of the PV flow rate in the motion-reconstruction cases with respect to the reference flow rate MAE = 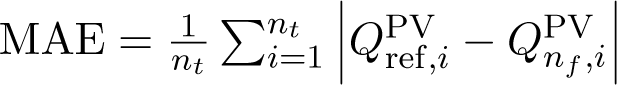, where n*_t_* is the number of time steps.

| $n_f$ | A | B | C | D | E | average |
| --- | --- | --- | --- | --- | --- | --- |
| 5 | 85.94 | 93.97 | 83.8 | 83.49 | 86.01 | 86.64 |
| 10 | 52.12 | 49.35 | 41.05 | 48.06 | 58.29 | 49.77 |
| 20 | 13.60 | 12.63 | 10.75 | 12.50 | 18.16 | 13.53 |
| 40 | 2.52 | 2.06 | 2.02 | 2.77 | 1.78 | 2.23 |

### 3.3 Impact of LA Wall-Motion Temporal Resolution on the Dynamics of Hemodynamic Indices

For subjects A and D, Figure 4 shows the time evolution of the spatially-averaged blood specific kinetic energy. Since the cycle-to-cycle variability is low, as can be seen in the insets of Figure 4, we focus only on the last simulated cycle.

**Figure 4:**
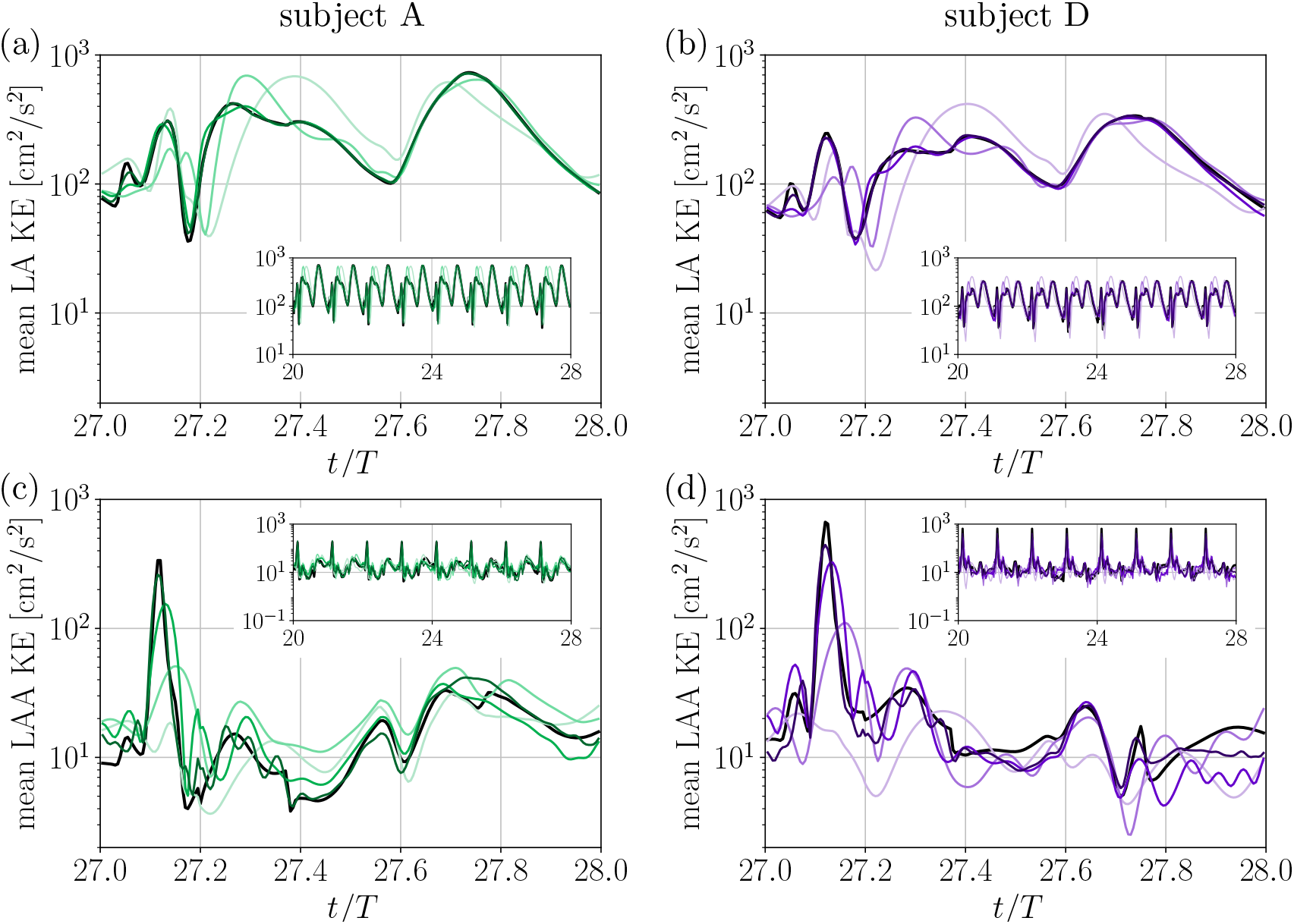
Spatially-averaged kinetic energy: Temporal evolution during one cycle in the LA of subjects (a) A and (b) D, and in the LAA of subjects (c) A and (d) D. The 5-, 10-, 20-, and 40-frame reconstructions are plotted in color scheme with an increasing number of frames from lighter to darker color, while the reference 4D LA reconstruction is plotted in black. The insets show the same quantity during 8 cycles.

Figure 5 presents the corresponding plots for the blood residence time. From the evolution of T*_R_* it is verified that the simulations indeed converged to a quasi-periodic state. However, cycle-to-cycle variability of T*_R_* is appreciable in the LAA. Together with the *T*_*R*_^LAA^ values being quite high, this justifies the decision to collect results over eight cycles after an initialization phase of 20 cycles.

**Figure 5:**
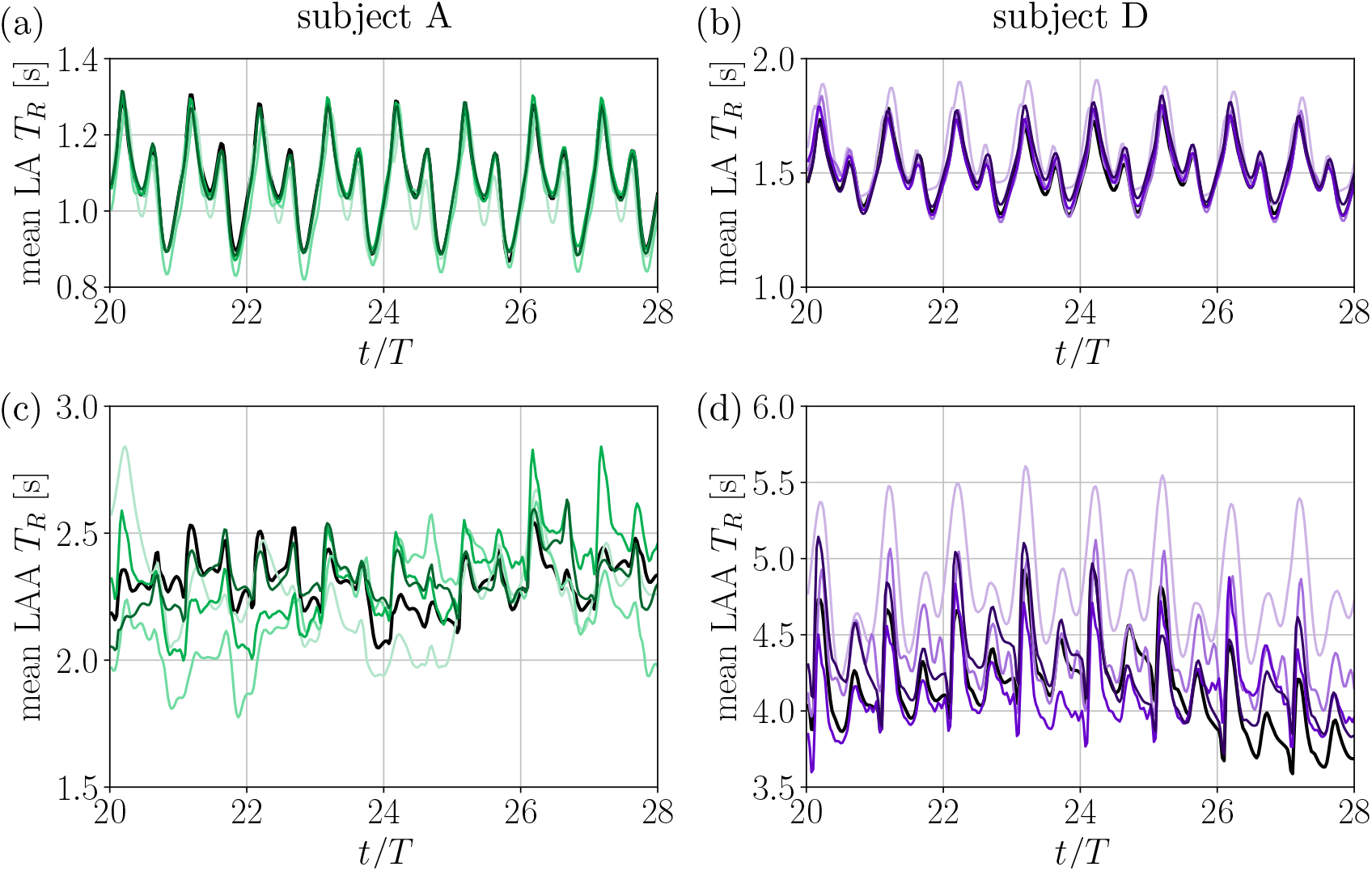
Spatially-averaged residence time: Temporal evolution in the LA of subjects (a) A and (b) D, and in the LAA of subjects (c) A and (d) D. Colors as in Figure 4.

In the LAA, the KE is systematically lower and T*_R_* consistently higher than in the LA, consistent with previous studies [16, 17, 20, 21, 31]. Regarding the impact of LA wall-motion reconstruction, we observe that CFD analysis based on the 5-and 10-frame reconstructions fail to faithfully reproduce the KE peak associated to the A-wave ( t/T *≈* 27.1) both in the LA and the LAA. Moreover, KE and T*_R_* in the LAA display similar spurious oscillations as those previously observed in the flow rate signals from the interpolated reconstructions in Figure 3.

### 3.4 Impact of LA Wall-Motion Temporal Resolution on Flow and Residence Time Fields

To gain further insight about the effect of LA wall motion, Figure 6 shows velocity vectors in a planar section and the 3D distribution of T*_R_* throughout the LA for subject D. Three time instants are considered: peak A-wave, an instant during the atrial expansion phase, and peak E-wave. For clarity, only regions with T*_R_ ≥* 2 s are shown and this quantity is entirely omitted for the snapshots of the atrial expansion phase.

**Figure 6:**
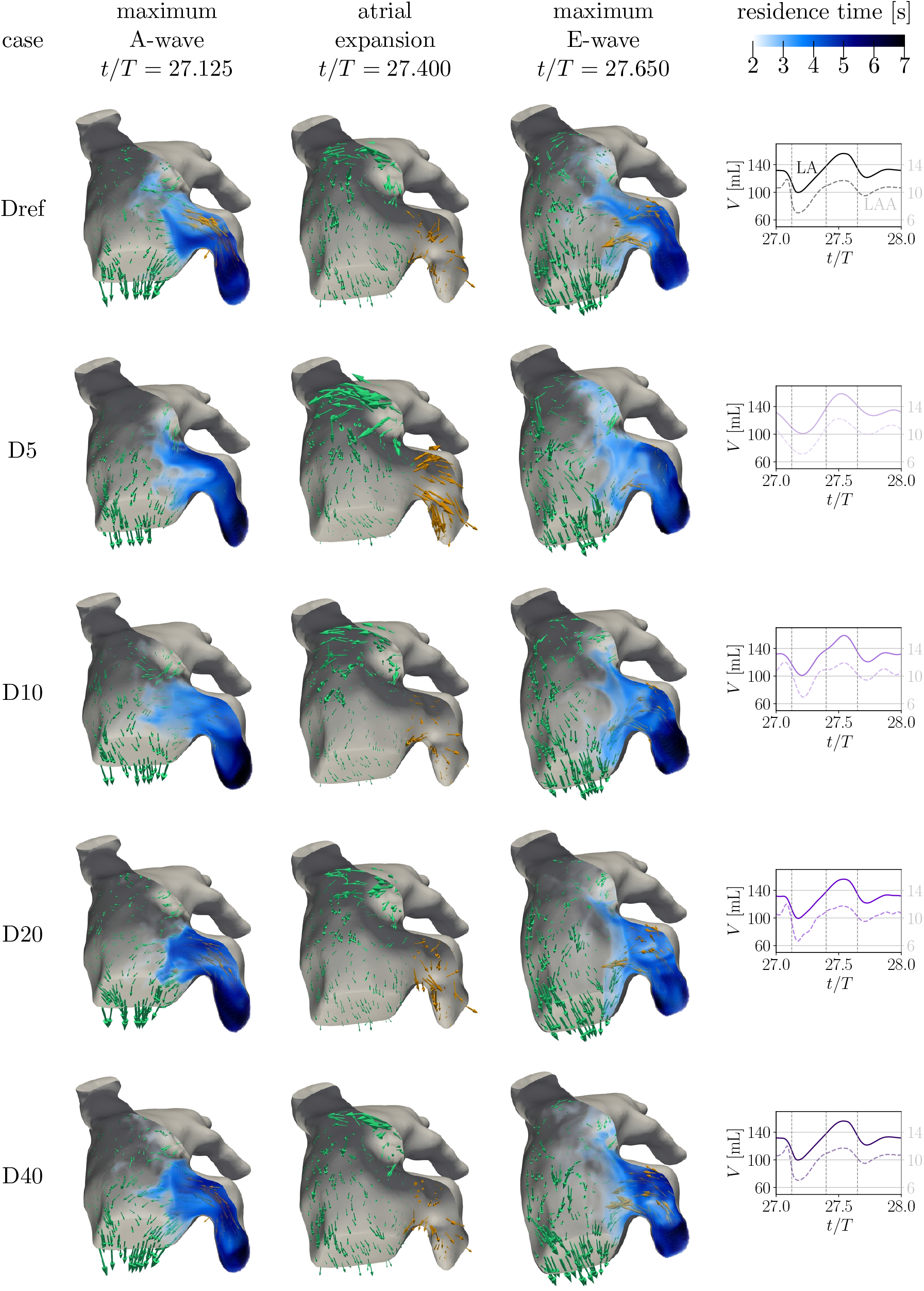
Visualization of the instantaneous residence time in the entire left atrium and velocity vectors in one planar section through the 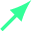 LA and 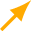 LAA of subject D at three time instants in the final simulated cycle, as indicated in the volume curves in the last column. During atrial expansion and E-wave, the LAA velocity vectors 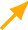 are elongated fivefold. For the time instant of atrial expansion, T*_R_* is omitted to facilitate visualization of the velocity vectors.

At peak A-wave, all simulations produce vigorous transmitral jets, given the identical imposed transmitral outflow rate. In addition, the simulations based on 20-and 40-frame 4D LA wall-motion reconstructions (D20 and D40) reproduce the LAA emptying flow observed in the reference simulation (Dref), whereas D5 and D10 exhibit only a weak outflow, suggesting that LAA emptying is degraded in these simulations. This is also verified by the evolution of LAA volume in the last column, which does not exhibit the steep decline observed for D20, D40, and Dref. Consequently, the mean T*_R_* in the LAA of D5 and D10 is elevated compared to the other cases (cf. Fig. 5d).

During the atrial expansion phase between the A-and E-waves, a pronounced flow is consistently observed near the LA roof, driven by PV inflow. In agreement with Q^PV^(t) depicted in Figure 3g,h, the inflow is most pronounced in D5 and weakest in D10 at this time instant, while in the other cases it is similar to the reference. All cases exhibit incoherent flow inside the LAA (yellow vectors), with the exception of D5, which experiences strong inflow.

At the E-wave peak, all cases appear similar because the LA functions primarily as a conduit, channeling freshly incoming blood from the PVs into the left ventricle (LV).

### 3.5 Impact of LA Wall-Motion Temporal Resolution on Flow Statistics

To quantify the effects described above while accounting for inter-subject variability, we analyze statistics for KE and T*_R_* inside each of the five subject’s LA and LAA.

#### Kinetic Energy Statistics

Figure 7a,b shows half-violin plots of the probability distributions of KE inside the LA and the LAA. As expected, the kinetic energy decreases from the LA to the LAA. Wall-motion resolution has only a minor effect on LA kinetic energy, with probability distributions that are nearly indistinguishable (Fig. 7a). The KE medians range between 39–130 cm^2^/s^2^ in the reference CFD simulations, and the simulations based on interpolated wall reconstructions slightly deviate from these values. On average over all five subjects, the medians of LA KE in the 5-, 10-, 20-, and 40-frame models deviate from their reference by 8.9%, 6.0%, 1.8%, and 1.4%, respectively.

**Figure 7:**
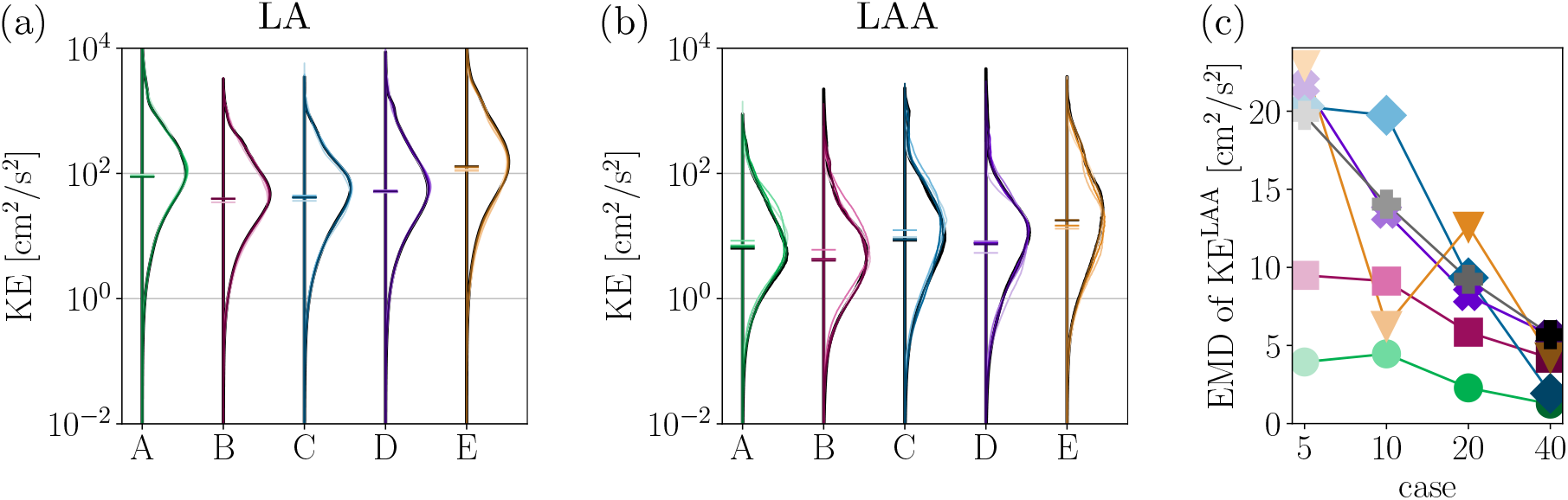
Kinetic energy: Probability distribution in the (a) LA and (b) LAA. The 5-, 10-, 20-, and 40-frame reconstructions of each subject are plotted in color scheme, with increasing number of frames from lighter to darker color, while the reference case is plotted in black for each subject. The horizontal lines indicate the medians. (c) Earth mover’s distance between the probability distributions of the reconstructed cases and their reference case for KE in the LAA of subjects 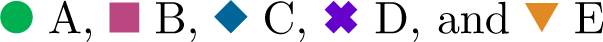. ✚ corresponds to results presented in §A.1.

In the 5-and 10-frame reconstruction cases, the fidelity of wall-motion reconstruction affects the LAA KE more noticeably (Fig. 7b), while no substantial differences are observed in the 20-and 40-frame cases. The LAA KE medians range from 4.4–17.7 cm^2^/s^2^, deviating by approximately 16%, 31%, 7%, and 5% from the reference, respectively, depending on the number of retained frames. For a more quantitative analysis, we computed the earth mover’s distance (EMD), which measures the similarity between two probability distributions [84]. For one-dimensional distributions, it can be computed as EMD 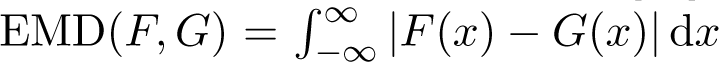 where F and G are the cumulative distribution functions [85]. In Figure 7c, we report the EMD between the reconstructed cases and their reference for KE in the LAA. Consistent with previous observations, discrepancies increase as n*_f_* decreases.

#### Residence Time Statistics

Similarly to the flow kinetic energy, T*_R_* is more sensitive to wall-motion reconstruction in the LAA than in the LA (Fig. 8). Across all patients, the global T*_R_* distribution is consistently bimodal (Fig. 8a), with a T*_R_ ≈* 0 s mode reflecting the LA’s reservoir role as fresh fluid enters during the atrial expansion, and a primary mode at nonzero times.

**Figure 8:**
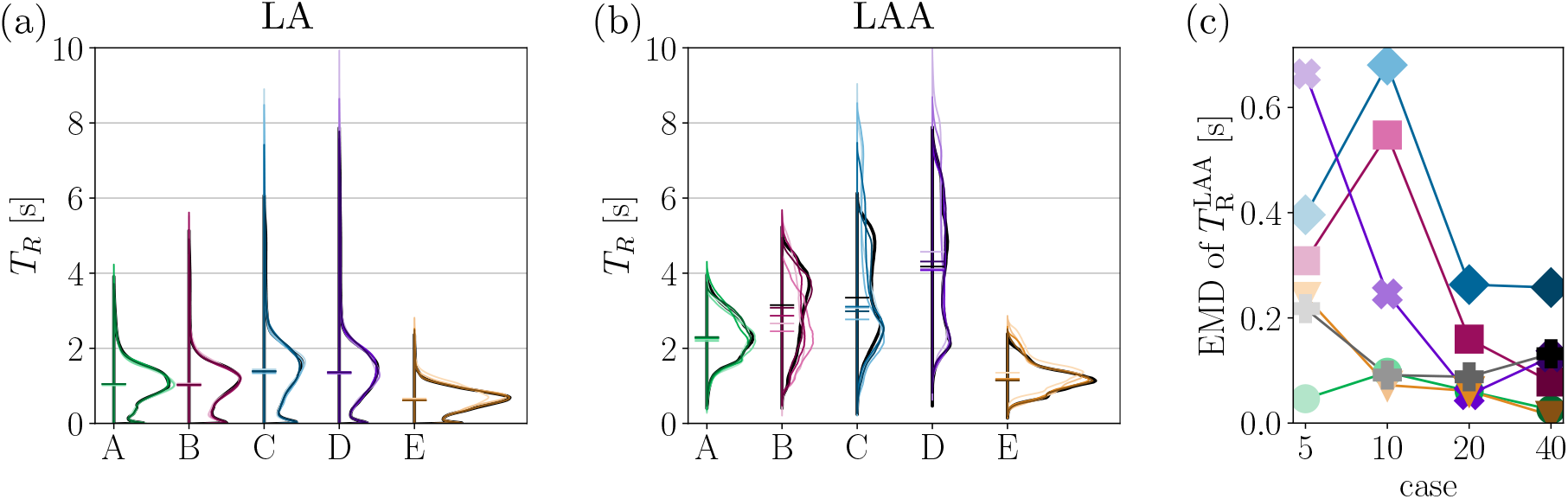
Residence time: Probability distribution in the (a) LA and (b) LAA. The horizontal lines indicate the medians. (c) Earth mover’s distance between the probability distributions of the reconstructed cases and their reference case for T*_R_* in the LAA. Colors and markers as in Figure 7.

Compared to the LA, the LAA experiences higher T*_R_* with greater inter-patient variability, consistent with its complex morphology. The distribution of T*_R_* in the entire LAA (Fig. 8b) is bimodal for some subjects (B, C, and D), an effect also observed by Dueñas-Pamplona *et al.* [16]. While the mode at a lower value mainly corresponds to fluid close to the LAA orifice, the second mode reflects fluid in its apex (not shown).

For the LA as a whole (median range: 0.61–1.39 s), the effect of motion reconstruction is minor, with medians deviating by 3.6%, 2.7%, 0.9%, and 0.4% from their reference, respectively. However, within the LAA, the differences are more pronounced: the T*_R_* medians span from 1.1 s to 4.2 s, deviating by 10.5%, 9.6%, 4.5%, and 3.5% from the reference values. The EMD for LAA T*_R_* (Fig. 8c) exhibits a similar trend.

#### Does Under-Resolved LA Motion Undermine CFD Stratification?

It may be acceptable to use patient-specific CFD based on standardized, under-resolved LA wall reconstructions for clinical decision support—even if the hemodynamic metrics are biased— provided the approximations preserve patient ranking (i.e., serve as surrogate indices for risk stratification). In that setting, the key requirement is to reproduce the ordering of patients by the metric, not its absolute values, which is a less stringent criterion than numerical accuracy.

To investigate this question, we present the median of KE and the 75^th^ percentile of T*_R_* inside the LAA in Figure 9, plotted vs. the number of frames retained in the LA wall reconstruction. As in prior sections, the median LAA KE generally departs from the reference value as the number of reconstruction frames decreases, with some exceptions at n*_f_* = 5. Even so, the rank ordering is largely preserved: subject E always exhibits the highest value, B is located at the lowest value, while A, C, and D fall in between. Likewise, the 75^th^ percentile of LAA T*_R_* shows irregular sensitivity to the number of retained frames. In general, subjects with lower T*_R_* exhibit less variations across the reconstructed cases. However, the ranking is preserved for all n*_f_*.

**Figure 9:**
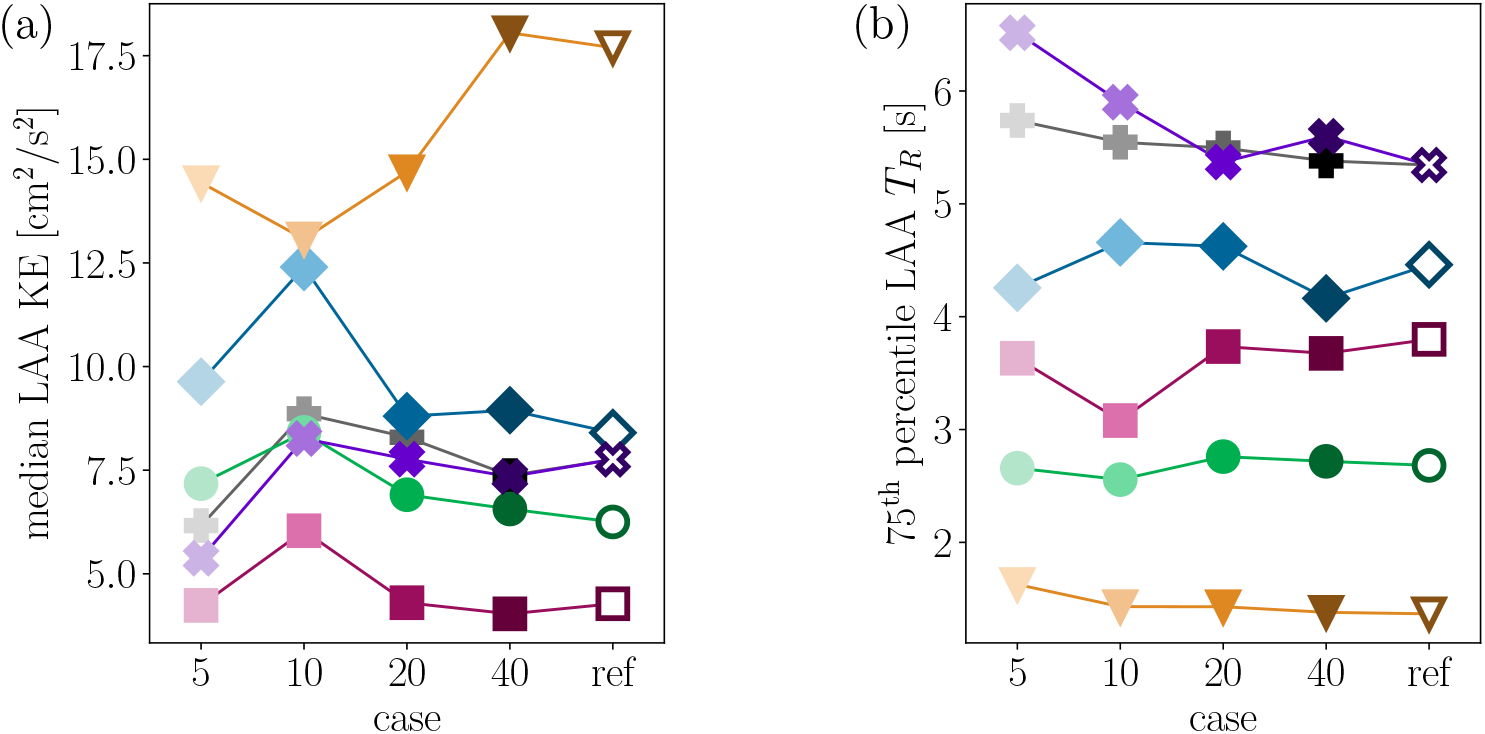
Hemodynamic metrics within the LAA of subjects 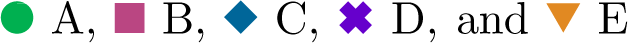. ✚ corresponds to results presented in §A.1. (a) Median kinetic energy and (b) 75^th^ percentile of the residence time.

## 4 Discussion

Left atrial (LA) wall motion changes rapidly as the chamber cycles through reservoir, conduit, and booster-pump phases each heartbeat. This imprints short-timescale features on filling and emptying flows via atrial volume changes and coupling with pulmonary venous (PV) inflow and left ventricle (LV) suction [86]. In patient-specific computational fluid dynamics (CFD), LA wall motion is reconstructed from 4D images with limited temporal sampling, which can blur these dynamics and introduce interpolation artifacts that propagate through mass balance into the boundary conditions and, ultimately, the simulated flow [32]. Critically, such errors persist even with fine spatial meshes and small solver time steps, because a numerically well-resolved CFD solver will faithfully propagate inaccuracies in the prescribed boundary conditions.

For robust clinical decision support, one would need to quantify how the temporal resolution of LA motion reconstruction influences metrics such as kinetic energy (KE) and residence time (T*_R_*), and whether coarser reconstructions still preserve patient ordering for risk stratification [87]. However, high-frame-rate imaging is often unavailable for technical, comfort, or safety reasons, limiting the accessibility to ground-truth LA wall reconstructions [88, 89]. Consequently, it remains an open question to which extent temporal undersampling distorts clinically relevant metrics—and when coarse reconstructions are still adequate for ranking.

In atrial fibrillation (AF), the loss of coordinated atrial systole and remodeling of reservoir and conduit dynamics alter the timing and amplitude of atrial function in complex, patient-specific ways [86], underscoring the importance of how LA wall motion is represented. Moreover, cardiogenic embolic events—including those strongly suspected to originate from the atrium— can occur even during sinus rhythm for large groups of patients [90–93]. Notably, embolic strokes of undetermined source (ESUS) occurring in the absence of AF may be mediated by LA cardiopathy. Although the underlying mechanisms are not entirely clear, these events are more prevalent among patients with substantial LA fibrosis, which is associated with disturbances in atrial electrophysiology, biomechanics, and flow dynamics [46, 91, 94, 95]. Therefore, evaluating the impact of temporal undersampling in sinus rhythm, where wall motion is more vigorous than in AF and exerts a strong influence on LA flow [31, 32], is also essential.

This manuscript presents a new approach to analyze the impact of LA wall-motion resolution on CFD-predicted hemodynamic metrics, where patient-specific atrial electromechanics (EM) simulations [46, 60] are used to generate highly resolved data that is used as surrogate ground truth. For each patient, reference CFD simulations are performed using LA wall motion directly obtained from the EM simulations. The corresponding moving LA anatomies are then temporally undersampled using n*_f_* = 5, 10, 20, 40 frames to mimic the typical resolution of 4D medical imaging, and additional CFD simulations are conducted using the interpolated LA models. The interpolated-wall simulation results are then compared to the reference results, with emphasis on KE and T*_R_* in the left atrial appendage (LAA), the most common site of LA thrombosis.

A substantial part of our effort involved prescribing the reference transmitral outflow rate Q^MV^ from the EM simulations across all CFD runs. This strategy ensured that differences in the results arose solely from the temporal sampling and interpolation of LA wall motion. Such an approach assumes that Q^MV^ is independent of the anatomical medical images, reflecting CFD workflows where the inflow is prescribed from Doppler, 4D flow MRI measurements, or mathematical models [16, 27, 50, 52, 96].

From the temporal evolution of LA volume, we observed that atrial contraction is poorly reproduced when using a low number of frames (n*_f_ ≤* 10) for motion interpolation, as the contraction occurs over a short timescale that is not resolved by the frames.

Global hemodynamic indices for the entire LA chamber, such as median KE and T*_R_*, were relatively insensitive to n*_f_*, with errors of less than 9% with respect to the reference simulations when varying n*_f_* between 5 and 40. However, CFD simulations based on LA wall-motion reconstructions with n*_f_* = 20 and 40 yielded favorable agreement, with errors averaging < 2% across our N = 5 patients. The observed range of variation was consistent with these quantities being mostly determined by the heart rate, mean LA volume, and LV stroke volume [17], which are relatively well captured even with n*_f_* = 5.

Focusing on LAA stasis, we found that deviations attributable to n*_f_* were minor compared to other effects. Larger variations in LAA stasis indicators arise from treating blood rheology as Newtonian versus non-Newtonian [17, 21], or from plausible physiological or anatomical changes, such as varying the PV inflow split ratios [20, 21] or PV orientation [27]. The n*_f_ ≤* 10 cases, however, occasionally fall within these physiological variability ranges. For all n*_f_*, the resulting variations in hemodynamic metrics remained smaller than those typically observed between moving-wall and fixed-wall simulations [21, 30–32]. Kjeldsberg *et al.* [32] further reported that the parametrized LA wall-motion model proposed by Corti *et al.* [37] improves the accuracy of hemodynamic metrics only when it is driven by patient-specific LA volume profiles with n*_f_* = 20, without examining other values of n*_f_*. This suggests that a coarse *patient-derived* LA wall-motion model may still be preferable to rigid walls, but the influence of temporal resolution in the LA volume profile remains unresolved.

An alternative, self-consistent method to reconstruct the LA wall geometry and prescribe Q^MV^ from a unique anatomical image acquisition is to segment time-resolved 4D sequences and use atrial surfaces for wall motion, while the time derivative of LV volume is used for transmitral flow [6, 17, 20, 31, 32, 49]. In these workflows, self-consistency implies that Q^MV^ is sensitive to image resolution as well. To assess how much this affects the CFD results, additional simulations were run, where the outflow rate was derived from undersampled LV volumes. Details are provided in the Appendix §A.1. Results showed that both KE and T*_R_* were largely unaffected.

Finally, we evaluated the effect of interpolating the underresolved wall motion with third-order splines [16, 22, 30, 32, 33, 51] in Appendix §A.2. Of note, spline interpolation reduced the spurious oscillations that arose in underresolved Fourier interpolants. However, the resulting CFD simulations produced similar hemodynamic metrics compared with the Fourier interpolation.

Taken together, these data suggest that patient-specific LA reconstructions derived from dynamic CT imaging provide a reliable basis for estimating LAA blood-stasis indices commonly used as surrogate markers of coagulation risk. Reconstructions with n*_f_ ≥* 20 exhibit the closest quantitative agreement with reference results, a frame rate achievable with most contemporary medical imaging modalities. When imaging data with n*_f_* < 20 must be used, the results deviate more noticeably from the ground truth. However, they may still provide approximate insights into LAA blood-stasis indices and related hemodynamic metrics. As n*_f_* decreases below 10, the patient ordering by KE or T*_R_* may not be preserved exactly, introducing potential inaccuracies near decision boundaries. Aggregating simulations based on different temporal resolutions may confound patient stratification. In such settings, standardized reconstruction protocols and explicit reporting of temporal resolution and interpolation strategy become critical to ensure that comparisons across patients or cohorts remain meaningful. Without such standardization, differences may reflect methodological inconsistencies rather than physiological variability.

### Limitations

This study is subject to several limitations that should be addressed in future work. First, atrial EM simulations were used as surrogate ground truth for wall motion and transmitral flow. Although these models have been validated [97], checked for physiological behavior [60, 67], offer high spatiotemporal resolution [66], and in our case generated physiologically realistic results (see §3.1), they still rely on modeling assumptions, parameter tuning, and simplified tissue properties that may not fully capture patient-specific atrial mechanics.

The size of the cohort is small (N = 5), which is partly attributed to the computational and storage requirements of the fully-resolved reference simulations. Although the cohort is diverse in anatomical shape and volume (see Figure 1 and Table 1), all subjects were modeled in sinus rhythm with homogeneous electromechanical myocardial properties (i.e., without fibrotic regions) and identical heart rates.

The risk of embolic stroke in AF patients has been shown to be sensitive to intricate details of LAA shape [98, 99]. However, the baseline LA anatomical segmentations used to drive the EM simulations were derived from MRI, yielding spatially smoother LAA representations than those typically obtained from modalities such as 4D–CT. The interaction between temporal and spatial resolution in anatomical modeling should be explored in future work.

Finally, all simulations were performed in sinus rhythm with preserved atrial contraction. As discussed above, this rhythm is clinically relevant, as embolic events can occur during sinus rhythm, even in subjects with subclinical intermittent AF [90]. Nonetheless, our findings may not fully extrapolate to AF, where atrial systole is absent or severely blunted. Prior work suggests that capturing wall motion may be less critical for flow predictions in AF due to reduced atrial dynamics [31], so the impact of temporal undersampling on stasis metrics may differ, warranting future investigation.

## 5 Conclusion

This study evaluated how CFD results are affected by the time resolution of patient-specific LA geometry data and the consequent interpolation of wall motion, which is necessary when the LA motion is derived from medical images. The findings indicate that the results based on 20 and 40 images per cardiac cycle exhibit no significant differences and closely align with the reference results. Reconstructions based on 10 and 5 images per cycle show larger deviations from the reference, without a consistent direction of bias, though patient ranking remains largely unaffected. Overall, these results support patient-specific reconstruction from dynamic CT imaging as a reliable tool for hemodynamic prediction across a range that overlaps with the feasible temporal resolution of clinical acquisitions. Higher frame rates are recommended when accurate absolute quantification, rather than relative stratification, is required, particularly within the LAA.

## Acknowledgments

This study was supported by the US National Institutes of Health (NIH) under award numbers R01HL158667 (NA, PMB, CMA, JCA) and R01HL160024 (JCA), and by the Austrian Science Fund (FWF) [10.55776/P37063] (CMA).

The computational results of the electromechanical simulations have been achieved using the Austrian Scientific Computing (ASC) infrastructure.

## Appendix

### A.1 Impact of Transmitral Flow Profile Temporal Resolution

Thus far, we have deliberately isolated the influence of LA wall-motion resolution. In every CFD run, the mitral outflow Q^MV^ was taken from the temporally fully-resolved reference EM simulation, ensuring that any differences arise from how the wall motion is sampled and interpolated rather than from changes in transmitral flow. This analysis is directly relevant to CFD workflows that use patient-specific or generic Doppler measurements to prescribe Q^MV^ [16, 27, 50, 52, 96]. However, many pipelines derive patient-specific Q^MV^ profiles from the time derivative of LV volume obtained from medical images [6, 17, 20, 31, 32, 49, 55, 56, 59]. In those workflows, the temporal resolution of the 4D imaging sequences affects not only the LA wall motion but also the estimated Q^MV^. This sensitivity carries to the PV inflow profiles via Equation 2, making both the boundary conditions and the CFD outputs sensitive to undersampling.

To investigate the impact of LV wall-motion resolution on the CFD results, we sampled the ventricular volume V ^LV^ provided by the EM simulations at the same time instants retained for reconstructing the LA motion, and subsequently interpolated and differentiated it in time using Fourier series. The mitral outflow rate was then obtained as Q^MV^ = 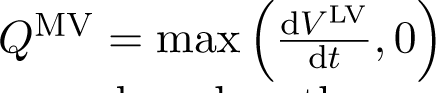, and new simulations were run for subject D. These additional simulations are based on the same LA wall motions as D5, D10, D20, and D40, respectively. However, to indicate the different treatment of Q^MV^, the cases are named with an asterisk, i.e. D5*, D10*, D20*, and D40*. The reference case Dref remains the same.

Figure 10a compares the transmitral flow rate profiles of cases D5*, D10*, D20*, and D40* with the reference profile. The E-wave (at t/T ≳ 0.55 in Dref) is captured reasonably well by all cases. The A-wave is considerably weakened and smeared in D5* and D10*, while it is accurately reproduced in D20* and D40*. Again, the Fourier interpolation leads to spurious oscillations in the reconstructed cases, especially for t/T ≳ 0.8.

**Figure 10:**
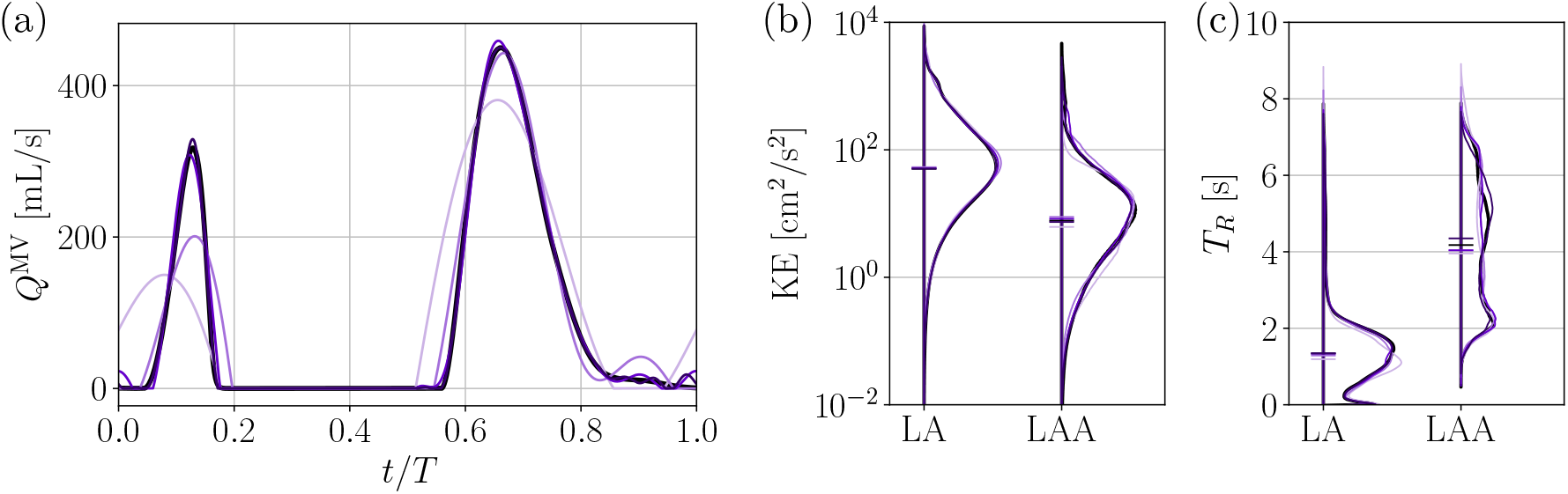
(a) Outflow rate through the mitral valve for case Dref, and the interpolated cases D5*, D10*, D20*, and D40*, where the flow rate is obtained from the time rate of change of LV volume as described above. Probability distribution of (b) KE and (c) T*_R_* for the cases of subject D. The respective regions of interest are indicated on the x-axis. The horizontal lines indicate the medians.

Figure 10b,c displays the half-violin plots for the results of KE and T*_R_* in the LA and LAA. The KE distributions of cases D5*, D10*, D20*, and D40* are similar to those from their corresponding cases D5, D10, D20, and D40, shown above (Fig. 7a,b), implying that the temporal resolution of transmitral flow profiles has a negligible incremental effect on the flow KE. Accordingly, the EMD and medians of LAA KE, which are displayed in gray markers in Figures 7c and 9a, follow a similar trend as when wall-motion is underresolved and inflow boundary conditions are fully resolved in time. While global T*_R_* is slightly underestimated by the 5-and 10-frame cases, they reproduce T*_R_* in the LAA more faithfully than for fully resolved inflow profiles (cf. Fig. 8b). This is also reflected in the EMD and 75^th^ percentiles of *T*_*R*_^LAA^ (see Figs. 8c and 9b), which are closer to their reference case.

### A.2 Spline Interpolation for LA Wall-Motion Reconstruction

In Figure 3e,f, the Fourier-based interpolation introduced oscillations in the temporal derivative of LA volume. These oscillations propagated into the PV inflow profiles (Fig. 3g,h) and imprinted on the temporal evolution of KE and T*_R_* in the LAA (Figs. 4c,d and 5c,d). Both Fourier-based interpolation [17, 20, 31, 49, 50, 52, 58] and spline interpolation [16, 22, 30, 32, 33, 51] have been used in previous studies to reconstruct underresolved wall motion. We therefore sought to assess whether the conclusions of the present study depend on the choice of temporal interpolation method. To this end, we repeated the simulations using a third-order spline interpolation with periodic boundary conditions instead of the Fourier-based approach. Specifically, for all subjects, we sampled the LA geometry at the same time instants as before (with n*_f_* = 5, 10, 20, 40), and prescribed the reference transmitral outflow rates Q^MV^ to all cases.

Figure 11 shows the resulting evolution of LA volume and its temporal derivative for subject D. As expected, the cubic splines reduced the spurious oscillations that arose in the underresolved Fourier interpolants. However, the underlying hemodynamic conclusions remain unchanged: the median kinetic energy and 75^th^ percentile of residence time in the LAA, reported in Figure 12, show variations very similar to those observed with the Fourier interpolation (Fig. 9).

**Figure 11:**
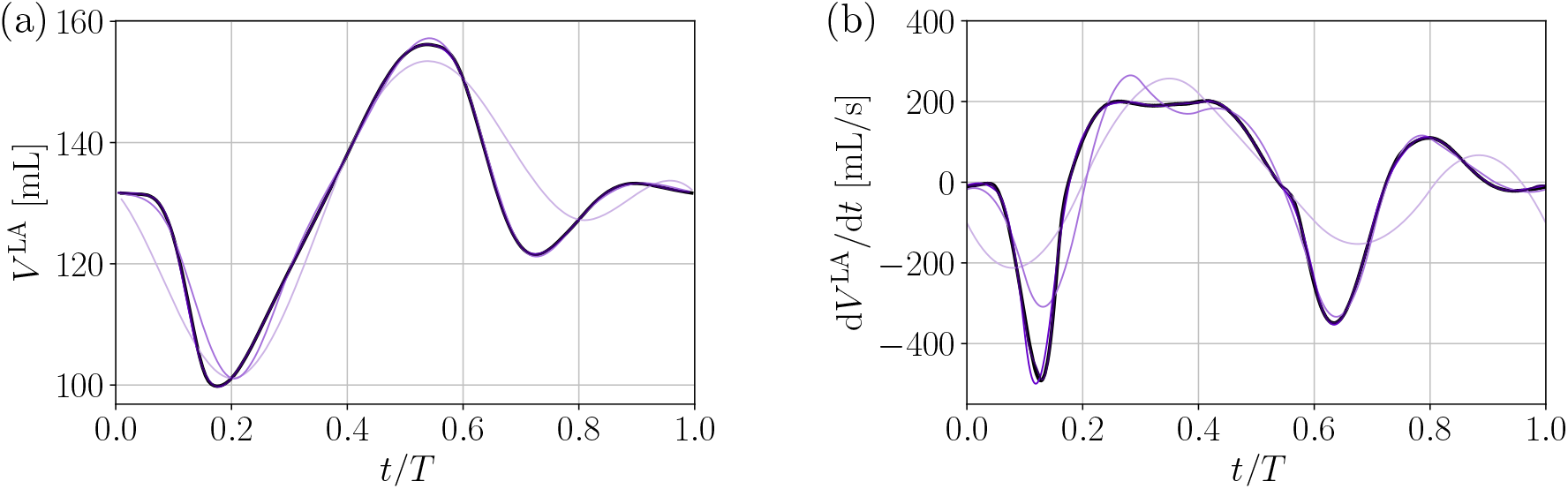
Temporal evolution of (a) LA volume and (b) time derivative of LA volume during one cycle for subject D. Motion reconstruction in the interpolation cases was performed with 3^rd^-order spline interpolation.

**Figure 12:**
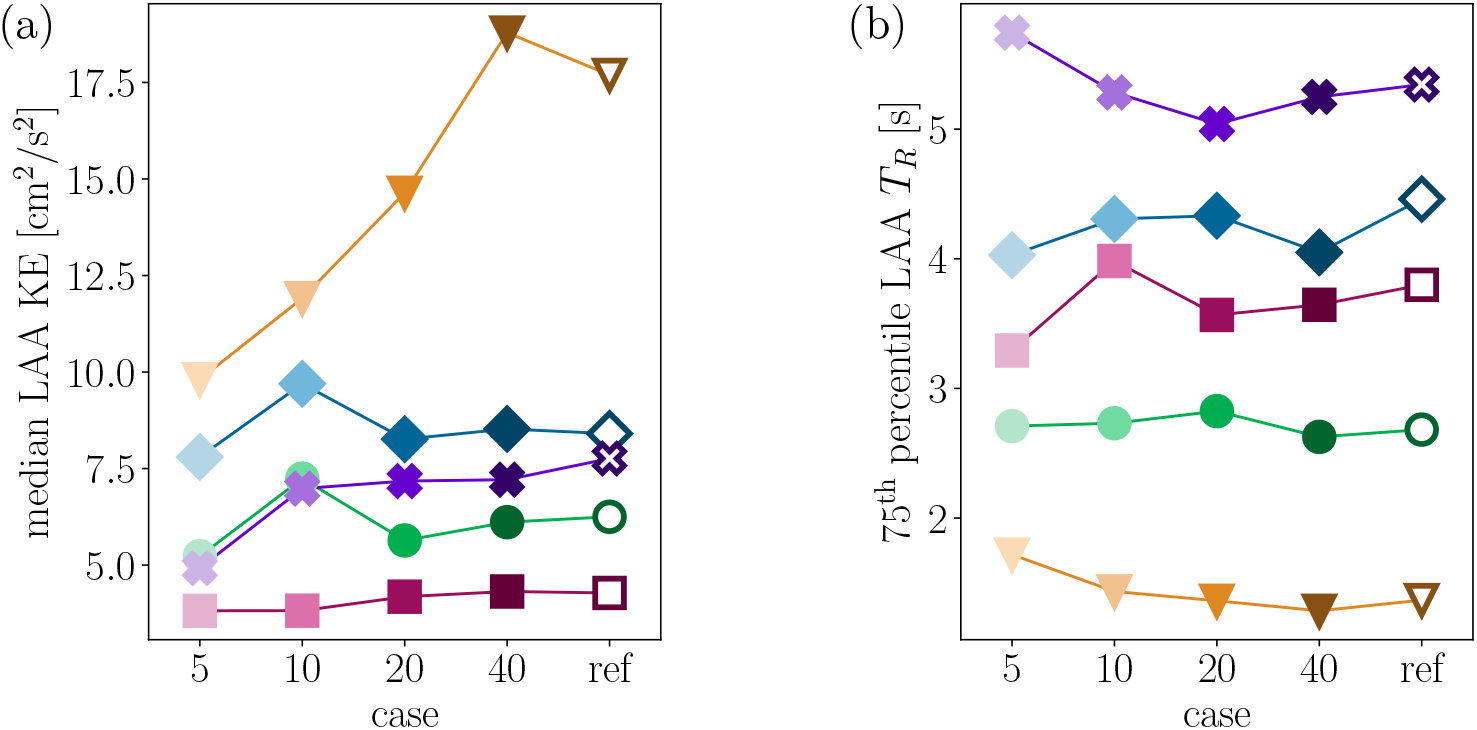
Hemodynamic metrics within the LAA of subjects 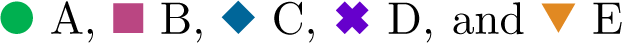 obtained by using 3^rd^-order spline interpolation for motion reconstruction. (a) Median kinetic energy and (b) 75^th^ percentile of the residence time.

## Acronyms

4D–CT: time-resolved computed tomography
AF: atrial fibrillation
CFD: computational fluid dynamics
CT: computed tomography
EM: electromechanics
EMD: earth mover’s distance
IBM: immersed boundary method
LA: left atrium
LAA: left atrial appendage
LV: left ventricle
MRI: magnetic resonance imaging
MV: mitral valve
PV: pulmonary vein

